# A National Estimate of the Health and Cost Burden of *Escherichia coli* Bacteraemia in the Hospital Setting: The Importance of Antibiotic Resistance

**DOI:** 10.1101/153775

**Authors:** NR Naylor, KB Pouwels, R Hope, N Green, KL Henderson, GM Knight, R Atun, JV Robotham, SR Deeny

## Abstract

**Background:** Antibiotic resistance poses a threat to public health and a burden to healthcare systems. *Escherichia coli* causes more bacteraemia cases in England than any other bacterial species, these infections, in part due to their high incidence, also pose a significant antibiotic resistance burden. The main aim of this study was to estimate the impact of *E. coli* bacteraemia on patient in-hospital mortality and length of stay. Secondarily, this study also aimed to estimate the effect of antibiotic resistance on these outcomes.

**Methods and Findings:** Case patients were adult *E. coli* bacteraemia patients infected between July 2011 and June 2012, as reported in an English national mandatory surveillance database, with susceptibility data taken from a national laboratory surveillance database. Control patients were all non-case, adult patients with an English hospital admission record. Case and control patient characteristics and admission information were taken from NHS Digital Datasets. ‘Resistance’ was defined as non-susceptible and intermediate isolates, whilst ‘susceptible’ was defined as susceptible and non-tested isolates. Time to in-hospital mortality and discharge was investigated through Cox proportional hazards models. To acquire estimates of excess length of stay, multistate models were constructed, with a unit bed day cost applied to estimate cost burden. The total number of case and control hospital spells was 14,051 and 8,919,275 respectively. Acquisition of *E. coli* bacteraemia was associated with a statistically significant increased daily risk of in-hospital mortality, especially for the first eight days of someone’s hospital admission [Hazard Ratio = 2.77 (95% confidence interval; 2.61-2.94)]. Antibiotic resistance did not seem to significantly increase this risk further, though did significantly reduce risk of experiencing a discharge event (dead or alive). *E.coli* bacteraemia was estimated to cost £14,340,900 over the study period (rounded to the nearest £100), with resistance associated with excess costs per infection of £220 - £420 dependent on resistance type, for those where a significant impact was found (rounded to the nearest £10).

**Conclusions:** *E. coli* bacteraemia places a significant burden on patient health and on the hospital sector in England. Resistance is an important factor on length of stay with regards to such infections.

## Introduction

Ensuring that safe, high quality treatments for infections are available for use is not only important to patients and clinicians, but essential for sustainable health systems. The rising incidence of severe infections, such as bacteraemia due to *Escherichia coli,* combined with resistances to treatment options becoming more common worldwide (such as resistance to ciprofloxacin and third generation cephalosporins), has been listed as a particular risk to human health by World Health Organisation, European centre for disease control and the US centre for disease control [1–3]. For example, the overall incidence of *E.coli* bacteraemia increased by 15.6% from 2010 to 2014 within England, with an increase in the absolute number of resistant isolates reported over the same time period [4]. *E. coli* is also the most common cause of community-onset bloodstream infections in the elderly in the United States [5]. As a result, there have been a number of national and international initiatives focusing on tackling infections with antibiotic resistance-related pathogens [2,6,7]. One example of such an initiative in the hospital setting is that of the English Department of Health’s financial reward scheme attempting to induce hospitals to reduce Gram-negative infections (such as *E. coli*) by half by 2020 [8]. In the United States, Congress has approved $160 million to be spent on surveillance and antibiotic stewardship initiatives to combat antibiotic resistance [9].

Despite the recent recognition of the threat posed by antibiotic resistance, and very recent investment, there is limited information as to the economic and health burden of antibiotic resistance [7,10]. Previous studies which have attempted to quantify the health and economic cost associated with *E. coli* or *Enterobacteriaceae* related bacteraemia, have been limited to quantifying the impact of third generation cephalosporin resistance with a limited sample of hospitals [11,12], or did not account for patient risk factors and co-morbidities [13]. Evidence from such studies suggests that third generation cephalosporin resistance is significantly associated with increased mortality for patients [11,12], however not much is known about the impact of resistance to other commonly used drugs used in treatment pathways, such as ciprofloxacin, gentamicin or piperacillin/tazobactam. Previous evidence on the hospital length of stay impact of resistance, in the case of *E. coli* or *Enterobacteriaceae*, indicates that cephalosporin resistance or extended-beta lactamase production is associated with increased length of stay [11,12,14].However, these studies have not been able to utilise national level samples and therefore may suffer from bias introduced through the recruitment of the hospitals and patients included in the sample, reducing generalizability of findings [10–12,14].

While economic modelling can extrapolate from limited existing estimates as to the potential long term impact of infection and resistance [7] there remains a need to produce robust estimates of the current health and economic burden of infection due to both resistance and sensitive organisms, in order to prioritise intervention strategies, and demonstrate the economic and health benefits arising from antibiotic stewardship, new antibiotics, vaccines or susceptibility testing [10]. This study aims firstly to estimate the impact of *E. coli* bacteraemia on patient in-hospital mortality and length of stay. It secondly aims to estimate the effect of antibiotic resistance on these outcomes and the costs to the English National Health Service due to such infections over the study period.

## Methods

### Study Design

A retrospective cohort study was performed in the English secondary care setting utilising national whole system datasets. Data cleaning and subsequent statistical analyses were performed in R version 3.3.3, utilising R-packages data.table, survival, etm, mvna, mstate and geepack, available at CRAN (available at http://cran.r-project.org) [15–19].

### Study Population and Outcomes

#### (i) Cases Cohort

Case patients were defined as those who had a record in the *E. coli* bacteraemia national mandatory surveillance database between July 1^st^ 2011 and June 30^th^ 2012 [13]. The Public Health England (PHE) mandatory *E. coli* bacteraemia surveillance database, for this period, was linked with the PHE voluntary susceptibility surveillance database (LabBase2) in a previously described analysis on 30-day all-cause mortality for *E. coli* bacteraemia patients [13]. This linked dataset held information on microbiological testing, including antibiotic susceptibility and timing of infection, for each *E. coli* bacteraemia case. In order to estimate additional length of stay and mortality attributable to infection, these linked data were further deterministically linked with a national hospital administrative database, ‘Hospital Episode Statistics’ (HES) [20], using patients’ NHS numbers (a unique patient identifier). Further information on the hospital trust was obtained through deterministic linkage with Estates Return Information Collection [21], using unique provider codes.

Antibiotic resistance, for the purpose of this paper included non-susceptible and intermediate isolates, as done previously with similar datasets [22]. Resistance was defined in the following terms:

1. Resistance to at least one of the following antibiotics ciprofloxacin, third generation cephalosporins (ceftazidime and/or cefotaxime), gentamicin, piperacillin/tazobactams and carbapenems (imipenem and/or meropenem).
2. Resistance to ciprofloxacin, third generation cephalosporins, gentamicin and piperacillin/tazobactams, individually.

Susceptibility testing was performed at a hospital level, whereby 95% of hospital laboratories use the European Committee on Antimicrobial Susceptibility Testing (EUCAST) methodology [22].

*E. coli* bacteraemia patients that had not been tested for each resistance category were grouped into the relevant susceptible category to enable maximum statistical power given our sample. Descriptive statistics related to the non-tested and susceptible groups were used to verify this assumption.

Therefore, throughout this paper ‘resistant cases’ refers to non-susceptible and intermediate isolates and ‘susceptible cases’ to susceptible isolates and non-tested isolates (as defined according to the laboratory data).

Onset of infection was defined utilising time of specimen as a proxy for time of infection [13]. Community-onset cases were defined as those for which date of specimen was within the first two days of hospital admission, if the specimen date was greater than two days post-admission the case was classified as hospital-onset. If community-onset cases had a previous hospital discharge 14 days prior to date of specimen, these were reclassified as community-onset hospital-associated. This 14-day definition was chosen as a bacteraemia episode length was classified as 14 days in our analysis, consistent with previous work utilising the same datasets [13].

#### (ii) Control Cohort

Control patients were defined as all patients who had been admitted to a hospital in England (according to HES [23]), with no recorded *E. coli* bacteraemia (according to the surveillance database [13]) during that hospital stay. These are referred to as “non-infected” controls throughout the rest of this study.

Case and control patients that had been in hospital longer than 45 days were artificially right-censored, as were deemed outliers for the analysis and to be in line with previous work using a similar methodology [12].

#### (iii) Outcomes of Interest

The primary outcomes of this study were daily risk of in-hospital mortality or discharge (as measured by hazard ratios and cumulative incidence functions) and excess length of hospital stay (as measured by hazard ratios and in excess days). Same day events (e.g. admission and discharge) were treated as being 0.5 days [24].

### Statistical Methods

Descriptive statistics were summarised using median and interquartile ranges for continuous variables, and proportions (represented by percentages) for categorical variables.

#### (i) In-Hospital Mortality

Cox proportional hazards models were constructed for in-hospital mortality adjusted hazard ratio estimates for *E. coli* bacteraemia. The baseline covariates included in all adjusted models were age, sex, a ‘modified Elixhauser comorbidity index’ [25] and hospital trust type [21]. Age and Elixhauser comorbidity index values were centred to their respective means.

To estimate the impact of resistance on the daily hazard of in-hospital mortality, the above baseline covariates plus as a time-dependent resistance categorical variable, coded 0 for no infection, 1 for non-resistant infections and 2 for resistant infections, were included as the independent variables.

The proportional hazards assumption was checked utilising plots of corresponding Schoenfeld residuals over time [17]. Slight deviations were found for the infection variable over time, therefore observations were grouped into time periods to account for this, utilising a step function approach, to produce time-dependent coefficients [17]. Therefore our analysis presents two hazard ratios for each independent variable for two different time periods (e.g. the first eight days and from eight days onwards).

Using the above methodology allows for the computation of time- and covariate-adjusted estimation of independent variable impacts the daily risk of in-hospital mortality [12,24].

Significance was determined by observing the 95% confidence intervals, utilising given standard errors from the models and assuming approximate normality.

Though cause-specific Cox proportional hazards models are useful for establishing mortality cause-specific hazard ratios in relation to covariates, they do not account for the competing risk of being discharged. To estimate the cumulative incidence of in-hospital mortality events over the 45-day period, given that patients could also be discharged within that time, a multistate model approach was used. The structure and further description of the multistate model can be found below [16,26].

#### (ii) Length of Stay

To estimate the impact of infection and resistance on length of stay, whilst adjusting for time dependency and other baseline covariates, the Cox proportional hazards models described above were also constructed in relation to a discharge (alive or dead) outcome, in place of the in-hospital mortality. However, these models do not easily translate into time-adjusted excess length of stay in days. Therefore, multistate models were constructed (theoretical schematic of included health states shown in Figure 1).

**Figure 1:**
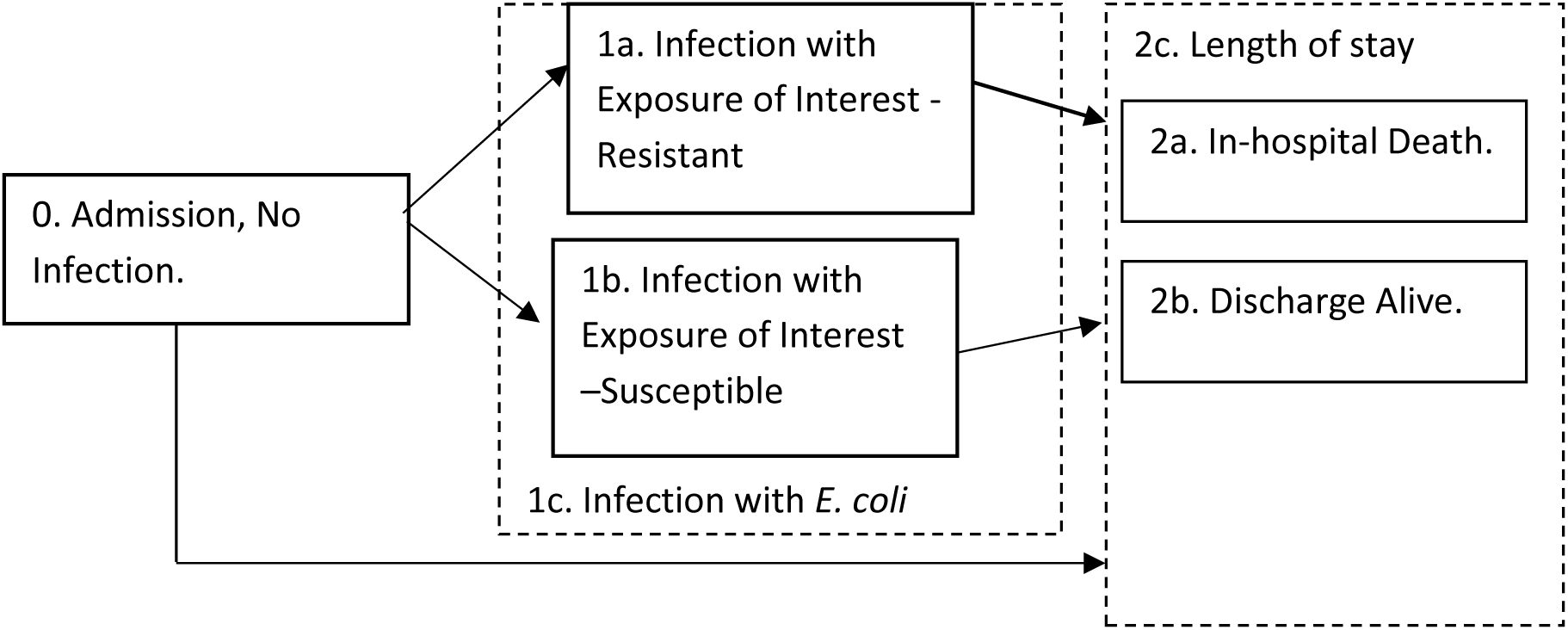
Multistate Model Schematic for Excess Length of Stay Estimation. The schematic represents the potential health states patients can travel through (in boxes) and the direction in which they may travel (via arrow directionality) in the multistate model. State 0 = “Admission, No Infection”, State 1 = “Infection of Interest” with 1a relating to resistant infections, 1b susceptible infections and composite state 1c being all *E. coli* infections, state 2a = “In-hospital Death” and state 2b = “Discharge Alive”. States 2a and 2b, together, form the length of stay endpoint, 2c.

The Aalen-Johansen estimator was used to estimate the time-varying transition probabilities between the health states within the multistate model, which utilises cumulative cause-specific hazard functions [12,27,28]. The expected length of stay (on each day) was calculated as a function of these transition probabilities [28]. The mean difference in estimated length of stay between transient states (depicted with 0 and 1.x notation in Figure 1) was calculated on each day (given that infection had or had not occurred by that day) and averaged across all days. This method for estimating transition probabilities is in line with previous studies utilising the multistate methodology to estimate infection-related, time-adjusted excess length of stay [12,14,24].

In practice, the built multistate model first compared general *E. coli* infection (1c, Figure 1) to admission (0, Figure 1), then subsequently compared resistant infections (1b, Figure 1) to susceptible infections (1c, Figure 1), whereby data were left-truncated to enable hospital onset patients to enter the model on getting the infection (i.e. enter at 1a or 1b at time of infection). The following equation was then solved;

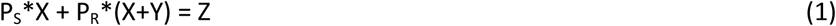

where P_S_ and P_R_ represent the proportion of susceptible and resistant cases respectively, Y represents the estimate for excess length of stay comparing resistant and susceptible cases, X represents the excess length of stay comparing susceptible cases to non-infected controls and Z represents the excess length of stay comparing all *E. coli* bacteraemia cases to non-infected controls.

In the multistate models standard errors were estimated through bootstrap sampling. The bootstrap sampling ran the models 1,000 times, randomly sampling through the transient state cohorts. This produced standard errors, which were utilised to estimate 95% confidence intervals, assuming approximate normality.

### Pseudo-observation analysis and estimation of confidence intervals

To adjust for time dependency and other covariates such as age, sex and comorbidity, pseudo-observations were created by running the multistate model, then running again after removing one person. The difference in excess length of stay estimates from the two model runs was then assigned to that person [16], noted as a pseudo-observation [12]. This was done for each person in the analysis.

The idea behind this method is that the ‘leave-one-out-diagnostic’ for the summary statistic (the length of stay estimate for the full sample minus one observation) contains information about the way in which covariates for each individual affect the estimator. The relationship between these pseudo observations and the covariates, including the exposure of interest, can subsequently be fitted using generalised estimating equations. Using this approach one can obtain length of stay estimates that are adjusted for confounders [29].

For feasibility reasons, a random sub-sample of 200,000 “non-infected” controls were selected and used for this (as opposed to the full “non-infected” control sample set).

95% confidence intervals were calculated to measure uncertainty within these models; these were defined assuming approximate normality and estimated using given standard errors.

#### (iii) Cost

Cost per spell was estimated by applying England’s Department of Health reference cost for an excess bed-day in secondary care to the number of excess days estimated [30]. This number was multiplied by the number of attributable spells (i.e. incidence of attributable hospital spells) to give an estimate of the annual burden in 2011-12. Results were rounded to the nearest £10 for cost per spell and to the nearest £100 for annual burden, given the lack of certainty surrounding the precision of such estimates. All costs presented are in 2012 Great British Pound (£).

95% confidence intervals were estimated by applying the unit cost to the 95% confidence interval lower and upper bounds for excess length of stay estimates. These bounds were then multiplied by incidence to estimate total annual cost 95% confidence intervals.

## Results

Across the period of interest (financial years 2011 – 2012 inclusive) 25,185,962 hospital episodes were initially retrieved from HES. 290,247 hospital episodes were linked to *E. coli* bacteraemia case patients. After applying the appropriate data cleaning to arrive at the desired case and control groups, 14,051 case and 8,919,275 control spells were included in this study. Tables 1A and 1B shows that the descriptive statistics of the cohorts analysed, with 1B focusing on within infection characteristics.

**Table 1:**
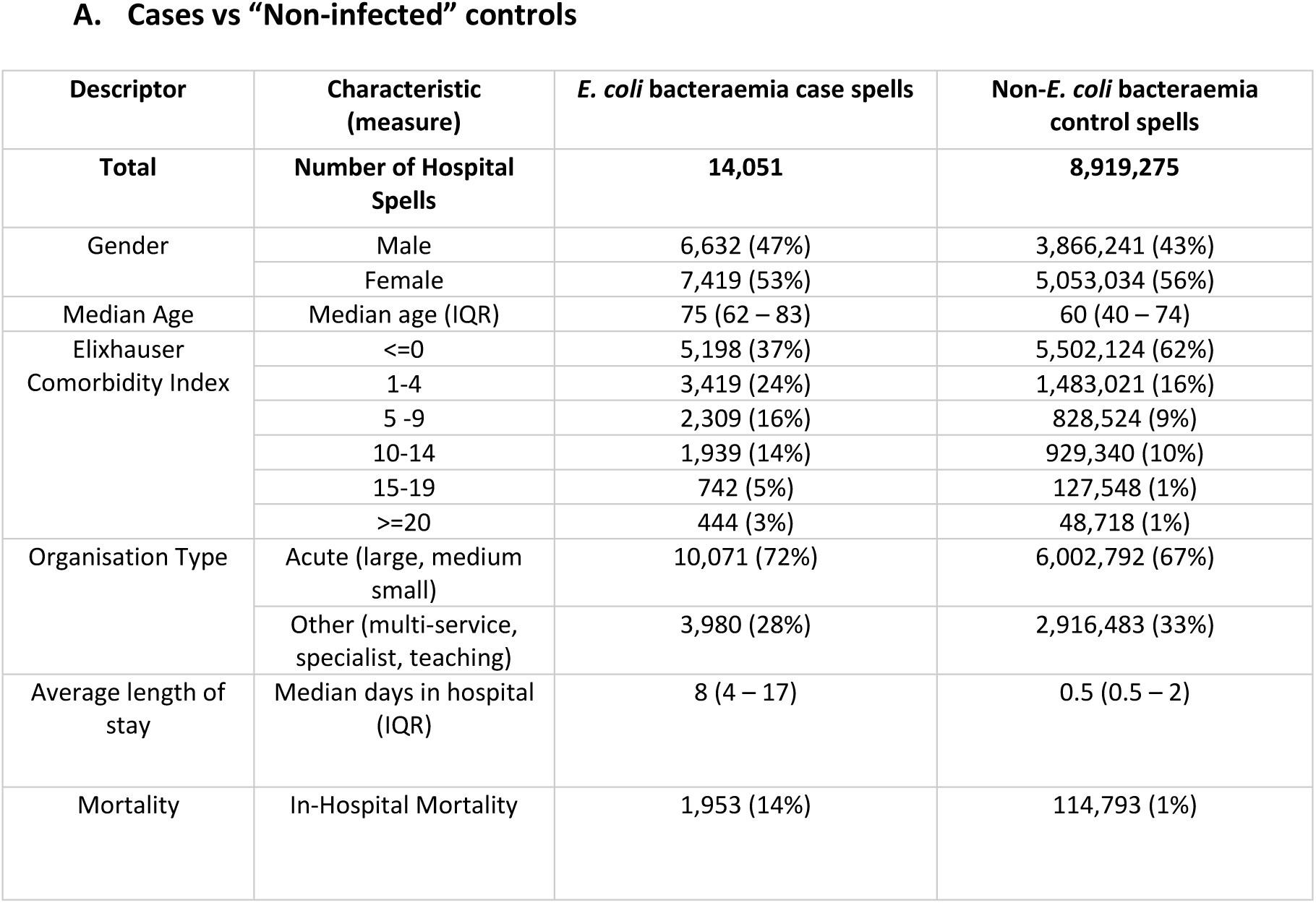
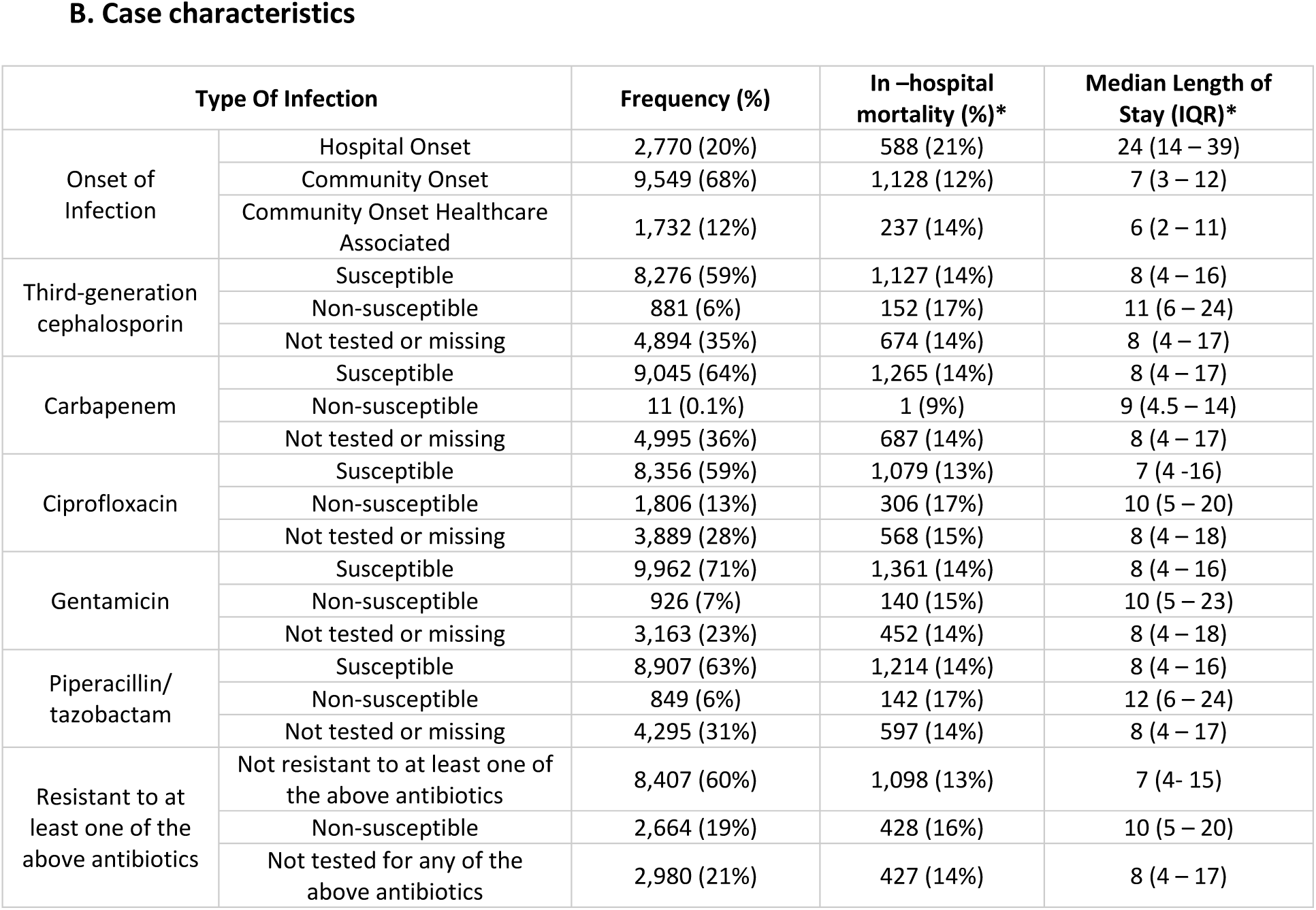
Descriptive Statistics for Case and Control Hospital Spells. Descriptive statistics are measured as a count (percentage) unless specified otherwise in the “Characteristic” column. *These are unadjusted descriptive statistics related to each row/characteristic group. Abbreviations: IQR; interquartile range

A. Cases vs "Non-infected" controls
| Descriptor | Characteristic (measure) | *E. coli* bacteraemia case spells | Non-*E. coli* bacteraemia control spells |
| --- | --- | --- | --- |
| Total | Number of Hospital Spells | 14,051 | 8,919,275 |
| Gender | Male | 6,632 (47%) | 3,866,241 (43%) |
|  | Female | 7,419 (53%) | 5,053,034 (56%) |
| Median Age | Median age (IQR) | 75 (62 – 83) | 60 (40 – 74) |
| Elixhauser Comorbidity Index | <=0 | 5,198 (37%) | 5,502,124 (62%) |
|  | 1-4 | 3,419 (24%) | 1,483,021 (16%) |
|  | 5 -9 | 2,309 (16%) | 828,524 (9%) |
|  | 10-14 | 1,939 (14%) | 929,340 (10%) |
|  | 15-19 | 742 (5%) | 127,548 (1%) |
|  | >=20 | 444 (3%) | 48,718 (1%) |
| Organisation Type | Acute (large, medium small) | 10,071 (72%) | 6,002,792 (67%) |
|  | Other (multi-service, specialist, teaching) | 3,980 (28%) | 2,916,483 (33%) |
| Average length of stay | Median days in hospital (IQR) | 8 (4 – 17) | 0.5 (0.5 – 2) |
| Mortality | In-Hospital Mortality | 1,953 (14%) | 114,793 (1%) |

B. Case characteristics
| Type Of Infection |  | Frequency (%) | In –hospital mortality (%)* | Median Length of Stay (IQR)* |
| --- | --- | --- | --- | --- |
| Onset of Infection | Hospital Onset | 2,770 (20%) | 588 (21%) | 24 (14 – 39) |
|  | Community Onset | 9,549 (68%) | 1,128 (12%) | 7 (3 – 12) |
|  | Community Onset Healthcare Associated | 1,732 (12%) | 237 (14%) | 6 (2 – 11) |
| Third-generation cephalosporin | Susceptible | 8,276 (59%) | 1,127 (14%) | 8 (4 – 16) |
|  | Non-susceptible | 881 (6%) | 152 (17%) | 11 (6 – 24) |
|  | Not tested or missing | 4,894 (35%) | 674 (14%) | 8 (4 – 17) |
| Carbapenem | Susceptible | 9,045 (64%) | 1,265 (14%) | 8 (4 – 17) |
|  | Non-susceptible | 11 (0.1%) | 1 (9%) | 9 (4.5 – 14) |
|  | Not tested or missing | 4,995 (36%) | 687 (14%) | 8 (4 – 17) |
| Ciprofloxacin | Susceptible | 8,356 (59%) | 1,079 (13%) | 7 (4 – 16) |
|  | Non-susceptible | 1,806 (13%) | 306 (17%) | 10 (5 – 20) |
|  | Not tested or missing | 3,889 (28%) | 568 (15%) | 8 (4 – 18) |
| Gentamicin | Susceptible | 9,962 (71%) | 1,361 (14%) | 8 (4 – 16) |
|  | Non-susceptible | 926 (7%) | 140 (15%) | 10 (5 – 23) |
|  | Not tested or missing | 3,163 (23%) | 452 (14%) | 8 (4 – 18) |
| Piperacillin/tazobactam | Susceptible | 8,907 (63%) | 1,214 (14%) | 8 (4 – 16) |
|  | Non-susceptible | 849 (6%) | 142 (17%) | 12 (6 – 24) |
|  | Not tested or missing | 4,295 (31%) | 597 (14%) | 8 (4 – 17) |
| Resistant to at least one of the above antibiotics | Not resistant to at least one of the above antibiotics | 8,407 (60%) | 1,098 (13%) | 7 (4 – 15) |
|  | Non-susceptible | 2,664 (19%) | 428 (16%) | 10 (5 – 20) |
|  | Not tested for any of the above antibiotics | 2,980 (21%) | 427 (14%) | 8 (4 – 17) |

### (i) In-hospital Mortality

From the adjusted Cox proportional hazards model, which adjusts for patient and hospital characteristics, it is shown that patients with an *E. coli* bacteraemia had around twice the daily risk of experiencing in-hospital mortality compared to those that did not have an *E. coli* bacteraemia. The impact of infection on mortality is higher during the first eight days of someone’s infection-related hospital stay [hazard ratio (HR) = 2.77 (95% confidence interval (CI); 2.61 – 2.94)] compared to after eight days [HR= 1.54 (95% CI; 1.43 – 1.65)].

Infection with a strain resistant to at least one of the tested antibiotics did result in a higher point estimate hazard ratio compared to susceptible strains, however this was not statistically significant (Table 2).

**Table 2:**
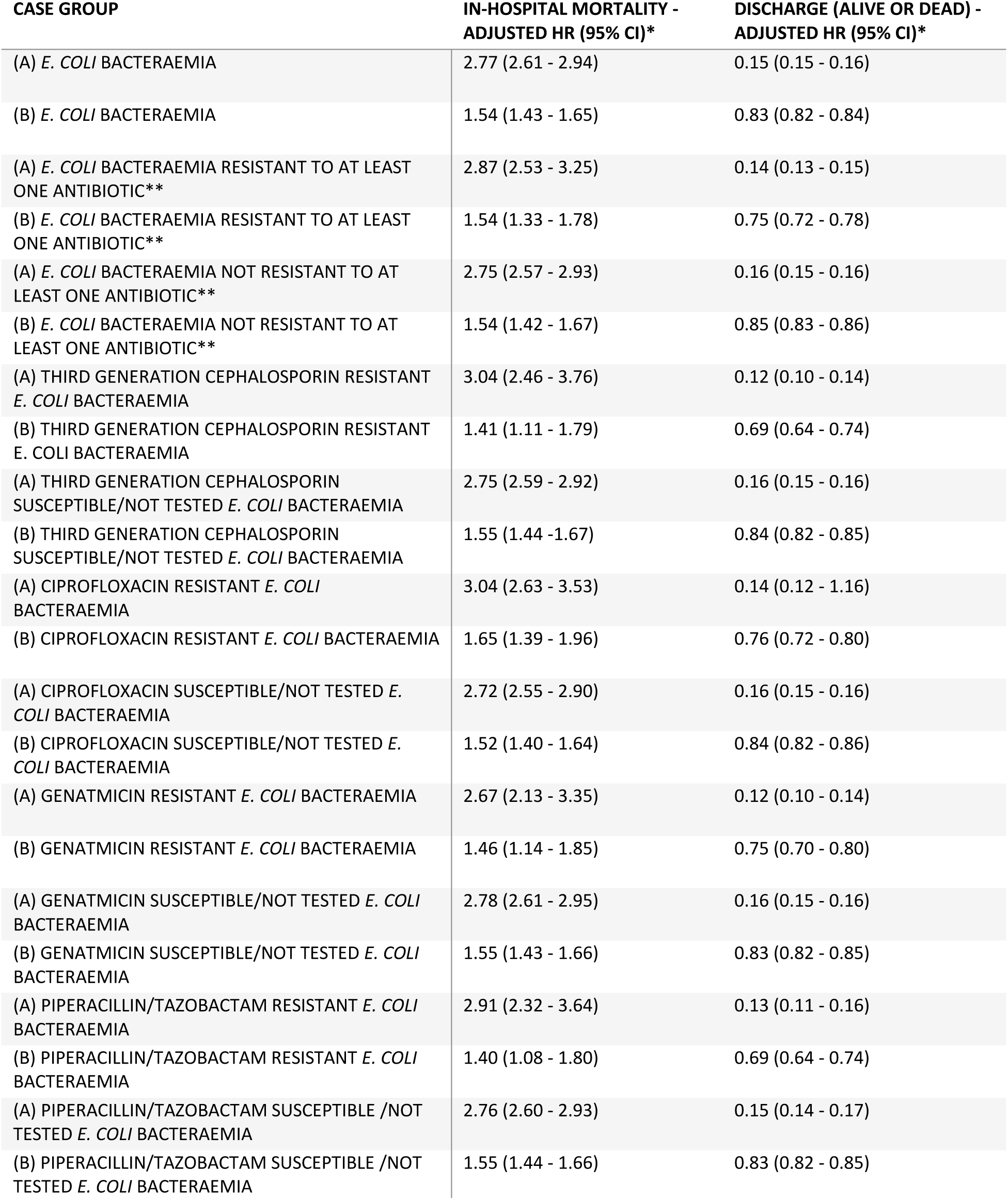
Daily Risk of In-Hospital Mortality of Discharge. Cause-specific hazard ratios for time to in-hospital mortality or discharge were estimated through Cox proportional hazards models, the comparator for all models was “non-infected” controls. Note that day zero refers to the day of admission. For in-hospital mortality (A) refers to the first eight days of admission and (B) refers to greater than eight days from admission. For discharge (A) refers to the first two days of admission and (B) refers to greater than two days from admission. *Adjusted for age, sex, Elixhauser comorbidity index and organisation type. **Tested antibiotics included ciprofloxacin, third generation cephalosporins, gentamicin, piperacillin/tazobactam and carbapenems. Abbreviations: CI; confidence interval, HR; hazard ratio.

Utilising a multistate model approach to estimate cumulative incidence, whilst accounting for competing risks, found that 1.3% of non-infected patients experience in-hospital mortality compared to 14.3% of *E. coli* bacteraemia patients. The estimates for cumulative incidence of in-hospital mortality for resistant (to at least one of the tested drugs) and susceptible strains were 16.1% and 13.4% respectively. For a breakdown by each resistance type see Appendix Table A.

### (ii) Length of Stay

The Cox proportional hazards models results (Table 2) show that, having accounted for patient characteristics and hospital characteristics on admission, an *E. coli* bacteraemia significantly decreases the daily risk of experiencing a discharge event (alive or dead). Resistance was found to also have a significant impact on experiencing discharge event (Table 2). Just focusing on those with a hospital stay of greater than two days from admission, third generation cephalosporin and piperacillin/tazobactam resistance had the largest impact on hazard of experiencing a discharge event, reducing the likelihood of experiencing such an event [HR=0.69 (95% CI; 0.64 - 0.74) for both]. Ciprofloxacin resistance had the lowest impact on hazard of experiencing a discharge event across both time groups (Table 2).

From the multi-state model it was estimated that *E. coli* bacteraemia is associated with 3.87 (95% CI; 3.69 – 4.04) excess hospital days (Table 3). With third generation cephalosporin and piperacillin/tazobactam resistance having the largest effect on length of stay, in comparison to the other tested antibiotics. Third generation cephalosporin was associated with 1.58 (95% CI; 0.84 – 2.31) excess days compared to equivalent third generation cephalosporin susceptible infections. Ciprofloxacin resistance was not significantly associated with excess length of stay in the time adjusted model.

**Table 3:** Excess Length of Stay Estimated by Multistate models. *Utilising multistate model analyses. **Adjusted for sex, age and Elixhauser – representing the estimate for a female of mean study sample age with mean Elixhauser comorbidity index value, utilising pseudo-observation analyses. ***Tested antibiotics included ciprofloxacin, third generation cephalosporins, gentamicin, piperacillin/tazobactam and carbapenems. Abbreviations: CI; confidence interval.

| Case Group | Control group | Excess Length of Stay in Days (95% CI) – Time Adjusted* | Excess Length of Stay in Days (95% CI) – Time & Covariate Adjusted ** |
| --- | --- | --- | --- |
| *E. coli* bacteraemia | Non-Infected Controls | 3.87 (3.69 - 4.04) | 2.98 (2.73 - 3.23) |
| *E. coli* bacteraemia resistant to at least one antibiotic*** | *E. coli* bacteraemia not found to be resistant at least one antibiotic*** | 0.84 (0.37 - 2.31) | 0.96 (0.35 - 1.56) |
| Ciprofloxacin resistant *E. coli* bacteraemia | Ciprofloxacin resistant *E. coli* bacteraemia | 0.46 (-0.11 - 1.03) | 0.96 (0.27 - 1.66) |
| Third generation cephalosporin resistant *E. coli* bacteraemia | Third generation cephalosporin resistant *E. coli* bacteraemia | 1.58 (0.84 – 2.31) | 1.78 (0.87 - 2.70) |
| Gentamicin resistant *E. coli* bacteraemia | Gentamicin resistant *E. coli* bacteraemia | 0.89 (0.20 - 1.58) | 0.71 (-0.20 - 1.63) |
| Piperacillin/tazobactam susceptible *E. coli* bacteraemia | Piperacillin/tazobactam susceptible *E. coli* bacteraemia | 1.23 (0.50 - 1.95) | 1.27 (0.31 - 2.23) |

Comparing all exposures of interest to a “non-infected” control group provides the estimates displayed in Figure 2 (see Appendix Table B for numerical estimates). This suggests that excess length of stay due to susceptible infections is between three and four days, whilst excess length of stay due to resistant infections is larger, but also more varied and uncertain (dependant on type of resistance).

**Figure 2:**
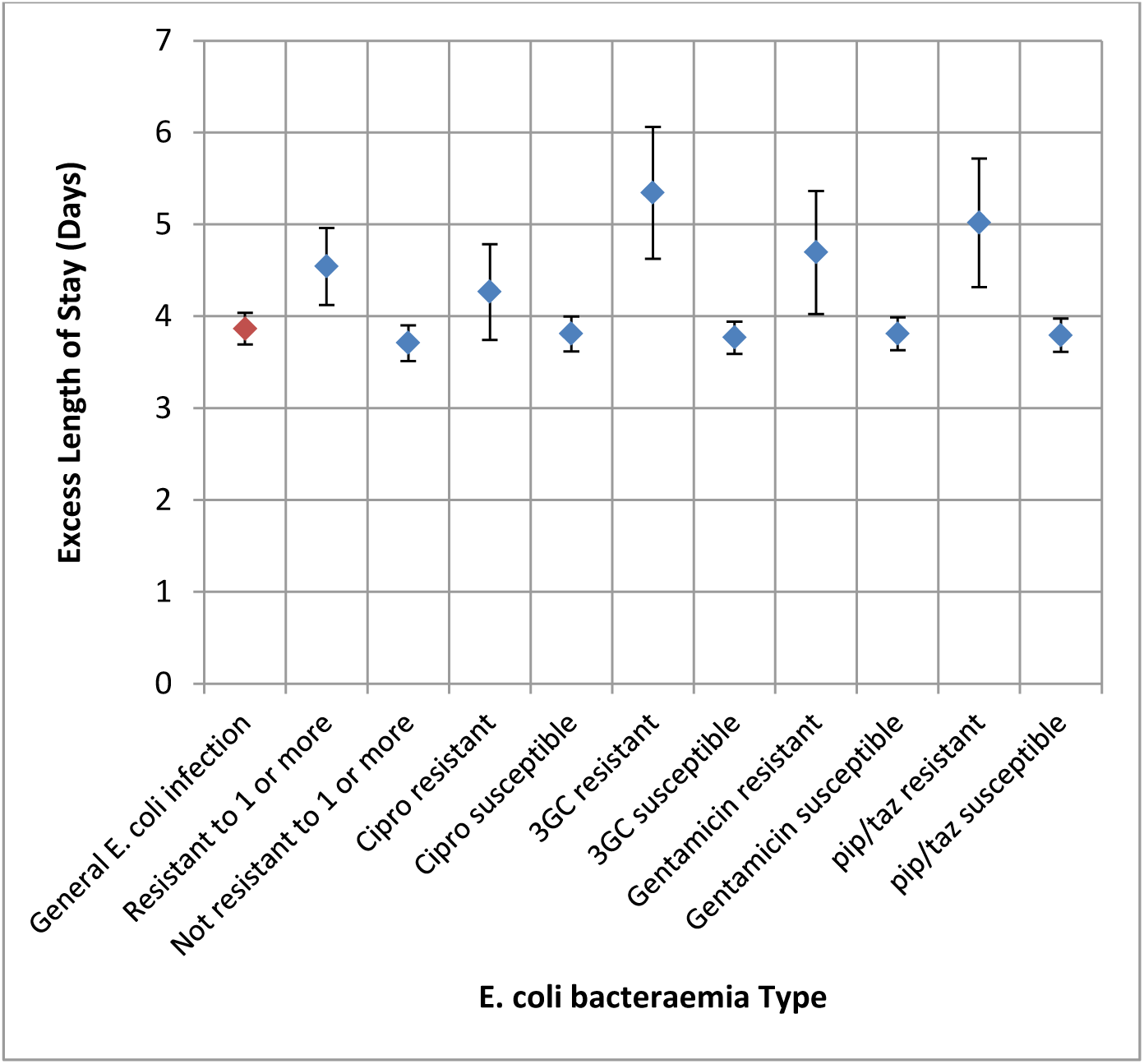
Excess Length of Stay of *E. coli* Bacteraemia as Estimated by Multistate Models & Algebraic Equations. Excess days associated with an exposure were calculated in comparison to “non-infected” controls. Multistate models were used to estimate *E. coli* bacteraemia versus “non-infected” controls estimate (red point). Other estimates (blue points) were derived utilising multistate model outputs in an algebraic equation [see equation (1)]. Error bars represent the 95% confidence intervals derived from bootstrapping. Abbreviations: 3GC; third-generation cephalosporin, *E. coli; Escherichia coli*, pip/taz; piperacillin/tazobactam. “Resistant to 1 or more” refers to being resistant to at least one of the tested antibiotics (ciprofloxacin, third generation cephalosporins, gentamicin, piperacillin/tazobactam and carbapenems).

The pseudo-observation analysis results presented in Table 3 represent estimates for a female patient of mean age and mean Elixhauser comorbidity of the subsample. For example, females 58 years old (mean age of the tested group), with a comorbidity index of 2 (mean comorbidity) with an *E. coli* infection are estimated to have 2.98 (95 % CI; 2.73 - 3.23) excess days in relation to patients with the same characteristics with no *E. coli* bacteraemia. Interestingly, adjusting for age and comorbidity (as described above) reduced the length of stay difference for general *E. coli* bacteraemia but increased the difference for specific resistance profiles, apart from for gentamicin resistance (Table 3). In this case, gentamicin resistance was not found to be significantly associated with excess length of stay.

### (iii) Cost

Applying the Department of Health Reference cost for an excess bed-day (£264 in 2011/12) [30] to the estimates for excess length of stay gives a cost per spell of per in-patient with *E. coli* bacteraemia of £1,020 (95% CI;£970 – £1,070). Utilising this cost per spell and number of spells, the estimated annual cost burden to hospitals due to *E. coli* bacteraemia in 2011/12 was £14,340,900 (Table 4). Adjusting solely for time dependency bias, excess annual costs associated with third generation cephalosporin resistance and piperacillin/tazobactam (compared to if they were susceptible infections) were £366,400 and £274,700 respectively (Table 4). That is to say, if all third generation cephalosporin resistant infections had been susceptible, over £350,000 would not have been spent on those infections (based on reduced length of stay).

**Table 4:** Excess costs associated with *E. coli* bacteraemia. Costs and 95% confidence intervals were derived applying a unit cost to estimates of excess length of stay. *Rounded to the nearest £10, ** Rounded to the nearest £100. Abbreviations: £; Great British Pound, CI; confidence interval. ***Tested antibiotics included ciprofloxacin, third generation cephalosporins, gentamicin, piperacillin/tazobactam and carbapenems.

| Infection | Incremental Excess Cost in 2012 £s (95% CI)* | Incidence per year for July 2011 – June 2012 (95% CI) | Excess Annual Cost in 2012 £s (95% CI)** |
| --- | --- | --- | --- |
| Compared to “non-infected” controls |  |  |  |
| *E. coli* infection | 1,020<br>(970 - 1,070) | 14051 | 14,340,900<br>(13,698,100 - 14,983,633) |
| Compared to susceptible cases |  |  |  |
| *E. coli* bacteraemia resistant to at least one antibiotic*** | 220<br>(100 - 340) | 2664 | 587,100<br>(258,400 - 915,900) |
| Ciprofloxacin resistant *E. coli* bacteraemia | 120<br>(-30 -270) | 1806 | 218,500<br>(-53,500 - 490,600) |
| Third Generation Cephalosporin resistant *E. coli* bacteraemia | 420<br>(220 - 610) | 881 | 366,400<br>(196,100 - 536,600) |
| Gentamicin resistant *E. coli* bacteraemia | 230<br>(50 - 420) | 926 | 217,000<br>(48,400 - 385,600) |
| Piperacillin/tazobactam susceptible *E. coli* bacteraemia | 320<br>(130 - 520) | 849 | 274,700<br>(111,900 - 437,400) |

## Discussion

Efforts to control antibiotic resistance, prevent infections and improve outcomes for patients have been hampered by a lack of investment, and uncertainty as to where to focus interventions; arising in part from the lack of robust evidence of the population health and health system costs associated with infection and resistance [7,10]. We found an excess cost of over £14 million associated with *E. coli* bacteraemia cases in adults in one year in, in English acute hospitals. Excess cost per resistant case was greatest in infections resistant to third generation cephalosporins; however this cost was found to be under £500 per resistant infection for all types of antibiotics. Infection with *E. coli* bacteraemia doubled patients’ probability of in-hospital death, though the impact of resistance was not statistically significant. Importantly, for both resistant and sensitive infections, excess mortality and delayed in discharge from hospital was significantly increased in the first eight days following infection.

### Comparison with previous findings

A study estimating the impact of third generation cephalosporin resistant and susceptible *Enterobacteriaceae* bloodstream infections in Europe found susceptible infections (compared to “non-infected” controls) to have an excess length of stay of 4.36 days (95 % CI; 3.91 – 4.81) [12]. This is similar to our estimate of 3.71 (95% CI; 3.29 - 4.13) (see Figure 2) for general *E. coli* bacteraemia. However, the impact of resistance in the European study was higher than in ours. We estimated resistance to third generation cephalosporins to increase length of stay (comparative to susceptible controls) by 1.58 (95% CI; 0.84 – 2.31) days, compared to their estimate of 3.53 days (95% CI; 2.08 – 4.96) [12]. However this could partly be due to the lower reliance on third generation cephalosporins in England compared to other antibiotics [31,32], though impact of all other antibiotic resistances (such as piperacillin/tazobactam) was still below 3 days in our analysis. This is the first study utilising this methodology to estimate the impact of other antibiotic resistances on length of stay (with regards to *E. coli* bacteraemia). Whilst just adjusting for time dependency in the multistate model, ciprofloxacin was found to not have a significant impact on length of stay. One reason for this could be the reduced usage of such fluoroquinolones in England, in an attempt to reduce *Clostridium difficile* infections [33].

In terms of impact on mortality, our Cox proportional hazards models found that once age, sex, comorbidity and trust type were adjusted for, resistance does not significantly impact mortality. In terms of cephalosporin resistance, this is not in agreement with a comparable European study on *Enterobacteriaceae*-related bacteraemia, which found resistance to have a significant impact [HR= 1.63; 95% CI: 1.13–2.35] [12]. However, it is in agreement with an earlier European study, which found such resistance in *E. coli* bacteraemia was not significantly associated with mortality [2.5 (95% CI; 0.9–6.8)] [11], though the cited study did likely suffer from power issues. A study on 30-day all-cause mortality utilising similar data-sources found that ciprofloxacin did have a significant impact on mortality [13]; however, this study was unable to adjust for patient comorbidities, which we did account for in our analysis.

Our findings suggest a number of clinical implications and priorities for future research and intervention. Firstly, as antibiotic resistant infections did add an excess cost to the health system, we would argue that *E. coli* infections, both antibiotic resistant and susceptible, are deserving of further research and investment. While there has been investment through initiatives to encourage infection control in the hospital setting [6–8], as we found that 68% of *E. coli* bacteraemia arose in the community, purely hospital focused initiatives may be limited in their impact. This would indicate that policies to improve the treatment of less serious infections (such as urinary tract infections) arising in the community that can develop into bacteraemia [22] could be something to be further investigated in terms of reducing the burden of *E. coli* bacteraemia. Furthermore, our findings indicate that for both resistance and susceptible infections the greatest impact on mortality can be achieved within the first eight days post admission. This is in line with current guidance around sepsis [32] which prioritise prompt identification of symptoms and treatment with appropriate antibiotics. To achieve similar outcomes for antibiotic resistant infections, while balancing the need for antibiotic stewardship, this will require further investment in the development of rapid diagnosis methods and implementation in the system [7].

### Strengths and limitations

This study is the first to estimate excess length of stay associated with antibiotic resistance bloodstream infections utilising a multistate model methodology and a nationally representative dataset, providing a large sample size of over 8 million control spells and over 14,000 case spells. It provides the first national estimate on the annual cost burden of *E. coli* bacteraemia. This study accounts for time-dependency bias and additional covariates in estimating excess length of stay, utilising novel techniques such as combining multistate models and generalized linear regression in pseudo-observation analysis. Both the cox models and multistate models are in agreement that piperacillin/tazobactam and third generation have the largest impact on time to discharge/excess length of stay, with ciprofloxacin estimated to have either the least or a non-significant impact. This reduces the likelihood that such conclusions are incorrect due to structural bias relating to the individual models.

One limitation of this study is that it is retrospective in nature and a cohort study based on data collected for other purposes, meaning that some data used may have been wrongly coded, missing or skewed. For example, the laboratory data collected on resistance profiles was from a voluntary database, meaning certain institutions with certain characteristics may more commonly report these isolate results. Our analysis did not include antibiotic exposure, since data were not available. It also did not explicitly compare site of infection onset (community as compared to hospital onset), though did account for timing of infection through time dependent covariates and multistate methodology [17,28]. The cut-off for inclusion in each category (2 days post-admission for inclusion in hospital onset) is arbitrary and may incorporate bias in its current form, as the appropriate control group for such analyses is unclear. In addition, there is no analysis in trends over time, as the scale of this analysis is limited to one year (July 2011 – June 2012). However, there has been little change in procedural policy on the treatment of such infections in England [34] meaning that the estimates are generalizable outside the years in the study cohort. Carbapenem resistance was not investigated individually, given that there were only 11 case spells in our sample, though was included as a potential resistance for the “resistant to one or more of the tested antibiotics” group. Carbapenem resistance is a major cause for concern [35], and investigation into the current and potential future impact of such resistance is needed.

The impact of resistance on mortality, length of stay and subsequent costs was estimated by grouping tested-susceptible and non-tested subjects. This may have introduced bias, as those not-tested could theoretically be resistant, however descriptive and time-to-event statistics were utilised to verify that this assumption. Bias introduced in this sense would mean our estimates are likely conservative. The costing does not account for any additional drug costs, procedural costs, ‘infection, prevention and control’ costs, or costs due to subsequent treatment in the community or readmission to hospital, meaning our estimates are likely conservative. However, recent literature suggests that length of stay is a key cost factor when estimating the costs of hospital onset infections [36], meaning our estimates likely account for a key proportion of costs. Our costing is an average unit cost of a hospital bed day and applied to all admission types, in reality a patient who was in intensive care could likely cost much more. Our estimates were rounded to nearest £10 and £100 for per case and annual burden respectively, this limits precision however was done as our methodology for costs meant that reporting costs to the nearest £1 could suggest we have produced estimates with misleadingly high granularity. Co-resistance was not taken into account within our analysis, since each infection type was analysed in the Cox and multistate models separately, to reduce computational complexity. Furthermore, in-hospital all-cause mortality is utilised as an outcome rather than total attributable mortality or 30-day mortality. Further data linkage for both cases and controls to the office of national statistics mortality dataset would allow the full impact of infection on mortality to be investigated in future research.

## Conclusions

The growing incidence of serious bacterial infections and the prevalence of antibiotic resistance are a threat to both patients and health systems. Our findings quantify the cost and mortality burden of *E. coli* bacteraemia and the influence of different resistances on this. Such findings will be useful for those identifying priorities for investment, infection control and modelling the health and economic impact of future trends in resistance. Additional research is therefore needed to identify modifiable health system and wider factors which could improve the outcomes of patients with resistant and susceptible *E. coli* bacteraemia, to aid the development of effective interventions aimed at reducing such infections. Our findings suggest such interventions could potentially improve care for patients while potentially reducing health system costs, though this is dependent on intervention effectiveness and costs.

## Acknowledgements

The authors would like to acknowledge Prof. Alan Johnson, Dr Berit Muller-Pebody, Mehdi Minaji and Esther Van Kleef, at Pubic Health England, for help in data access, management and coding.

## Appendix

**Table A:**
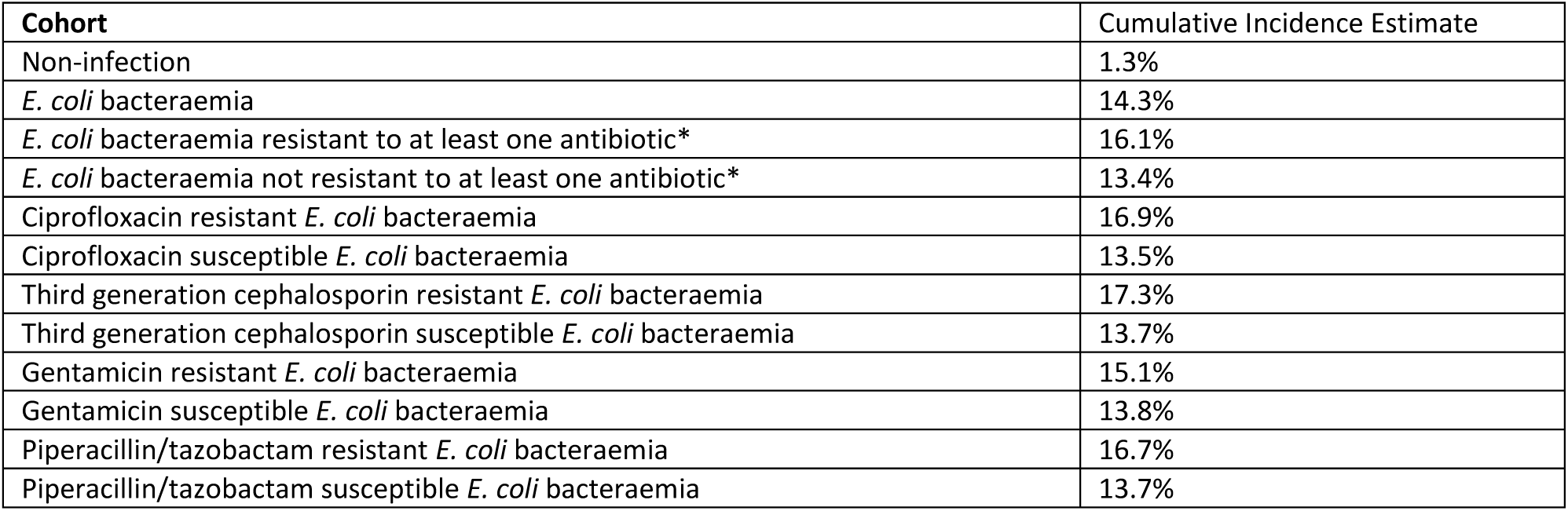
Cumulative Incidence of In-Hospital Mortality at 45 Days Post-Admission by Resistance Type. *Tested antibiotics included ciprofloxacin, third generation cephalosporins, gentamicin, piperacillin/tazobactam and carbapenems.

**Table B:**
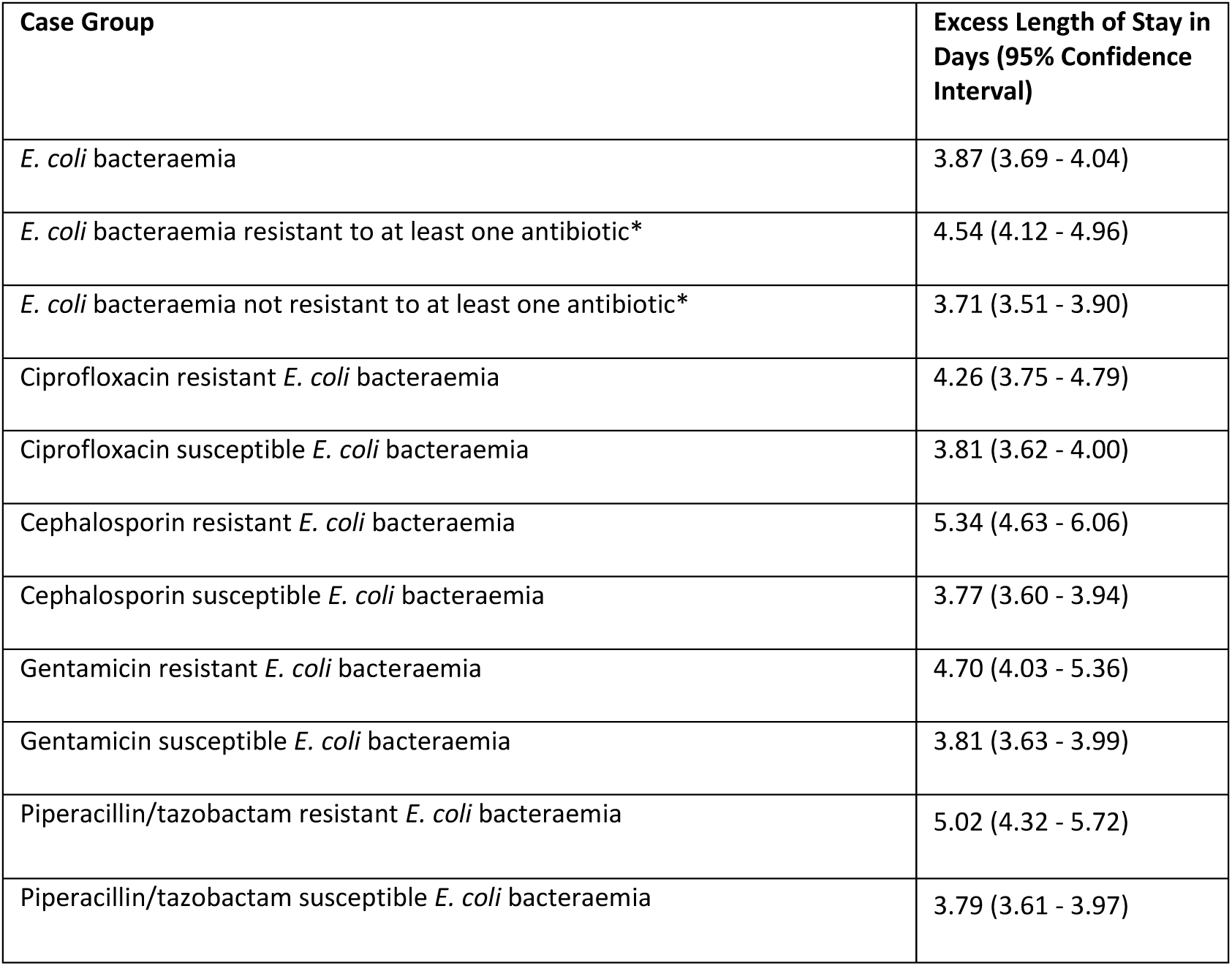
Excess Length of Stay in Comparison to “Non-Infected” Controls. *Tested antibiotics included ciprofloxacin, third generation cephalosporins, gentamicin, piperacillin/tazobactam and carbapenems.

